# Extensive lineage-specific rediploidisation masks shared whole genome duplication in the sturgeon-paddlefish ancestor

**DOI:** 10.1101/2022.05.16.492067

**Authors:** Anthony K. Redmond, Manu Kumar Gundappa, Daniel J. Macqueen, Aoife McLysaght

## Abstract

Whole genome duplication (WGD) is a dramatic evolutionary event generating many new genes and which may play a role in survival through mass extinctions. Paddlefish and sturgeon are sister lineages that both show genomic evidence for ancient WGD. Until now this has been interpreted as two independent WGD events due to a preponderance of duplicate genes with independent histories. Here we show that although there is indeed a plurality of apparently ‘independent’ gene duplications, these derive from a shared genome duplication event occurring close to the Permian-Triassic mass extinction period, followed by a prolonged process of reversion to stable diploid inheritance (rediploidisation). We show that the sharing of this WGD is masked by the fact that paddlefish and sturgeon lineage divergence occurred before rediploidisation had proceeded even half-way. Thus, for most genes the resolution to diploidy was lineage-specific. Because genes are only truly duplicated once diploid inheritance is established, the paddlefish and sturgeon genomes are a mosaic of shared and non-shared gene duplications resulting from a shared genome duplication event. This is the first time that lineage-specific resolution of genes from a common WGD event has been shown to affect such a large proportion of the genome.

## Introduction

Ancient WGD events have occurred across the tree of life and are especially well studied in plants^1–4^, yeast^5,6^, and vertebrates^7–18^. These events are often hypothesised to have facilitated evolutionary success through provision of the raw genetic materials for phenotypic innovation and species diversification^1,19–23^. A key evolutionary process after WGD is rediploidisation—the transition of a polyploid, usually tetraploid, genome to a more stable diploid state^1,7,9,14,22–25^. Importantly in this context, WGD events are derived from either hybridisation of two different parent species (allopolyploidisation) or from doubling of the same genome at the intra-species/individual level (autopolyploidisation), each with distinct cytogenetic outcomes. Classically, the non-homologous chromosomes of new allopolyploids preferentially pair bivalently during meiosis, whereas the four homologous chromosomes of new autopolyploids take a multivalent formation^1,9,14,22,24^. This results in ongoing homologous recombination, and hence gene conversion and homogenisation, across the four allelic copies at each locus. Suppression of recombination, probably achieved through chromosomal rearrangements and other mutations, is a necessary step to ‘rediploidise’ these genes into two distinct (bivalent) ohnolog loci (WGD-derived duplicate genes) from tetraploid alleles. It is only then that uninterrupted sequence and functional divergence can occur between ohnolog pairs^24^. The rediploidisation process thus uncouples the genome duplication process from the gene duplication process in autopolyploids, as it is only once rediploidisation has occurred that the locus can be considered duplicated.

Substantial evidence arising from studies of the ancestral salmonid WGD^18,24,26^, and to a lesser extent the ancestral teleost WGD^13,27,28^, indicates that autopolyploid rediploidisation can be temporally protracted, occurring asynchronously across the genome over tens of millions of years^13,18,24,26–28^. Major implications arise from the accompanying delay in any evolutionary processes that depend on ohnolog genetic divergence^24^. These include well-established models of functional evolution after gene duplication, e.g. sub-/neo-functionalization^29,30^, and models of reproductive isolation involving reciprocal loss of ohnologs in sister lineages^31^. Furthermore, if speciation occurs before rediploidisation has completed in descendent lineages of the same WGD, rediploidisation may occur independently in these daughter lineages (lineage-specific rediploidisation) in some genomic regions^24^. This in turn allows for ohnolog pairs to independently evolve divergent regulatory and functional trajectories in each lineage, potentially in response to lineage-specific selective pressures^24^. Ohnologs with this history have been described as following the LORe (**L**ineage-specific **O**hnolog **Re**solution) model^24^.

Although noted as a potentially important evolutionary process after several non-teleost WGDs^4,8,12,24,27,32–34^, it remains unclear whether asynchronous and lineage-specific rediploidisation is a general feature after autopolyploid WGD outside the teleost clade. The past two years have seen the generation of high-quality reference genomes from multiple non-teleost ray-finned fish lineages^7,8,35,36^, which share two ancient rounds of WGD common to all jawed vertebrates^9,11,37^, but lack the teleost-specific WGD event^7,8,35,36,38,39^. These species typically have slowing evolving genomes, enabling more reliable inference of the ancestral state of bony vertebrates than teleosts, while providing outgroups to understand the impact of the teleost-specific WGD event^7,8,35,36,39^. Among these newly available genomes are chromosome-scale assemblies for the sterlet sturgeon (*Acipenser ruthenus*)^7^ and American paddlefish (*Polyodon spathula*)^8^, which each have experienced WGD in their evolutionary histories.

Multiple WGD events have occurred during sturgeon evolution, but only one, thought to be shared by all sturgeons, is present in the sterlet sturgeon’s history^7,40–42^. On the other hand, American paddlefish is the only extant paddlefish^43^, and is suggested to have undergone a single WGD event^8,35,44–46^. Despite being sister lineages, together representing extant Acipenseriformes, previous analyses have consistently rejected a shared ancestral WGD in favour of independent WGD events (**Fig. 1**)^8,35,38,44,45^. Efforts to date these WGDs have produced incongruent results, with estimates ranging from 21.3 Ma^38^, 51 Ma^35^, and 180 Ma^7^ for the sturgeon WGD, and 41.7 Ma^44^, ∼50 Ma^8^, and 121 Ma^35^ for the paddlefish WGD. Although some authors have suggested that asynchronous rediploidisation may contribute to this incongruence^8,44^, this process has been ignored when dating these WGDs. Furthermore, the potential for genome-wide lineage-specific rediploidisation (e.g.^13,24,26^) to mask a shared WGD event has never been formally proposed or tested in any lineage.

**Figure 1.**
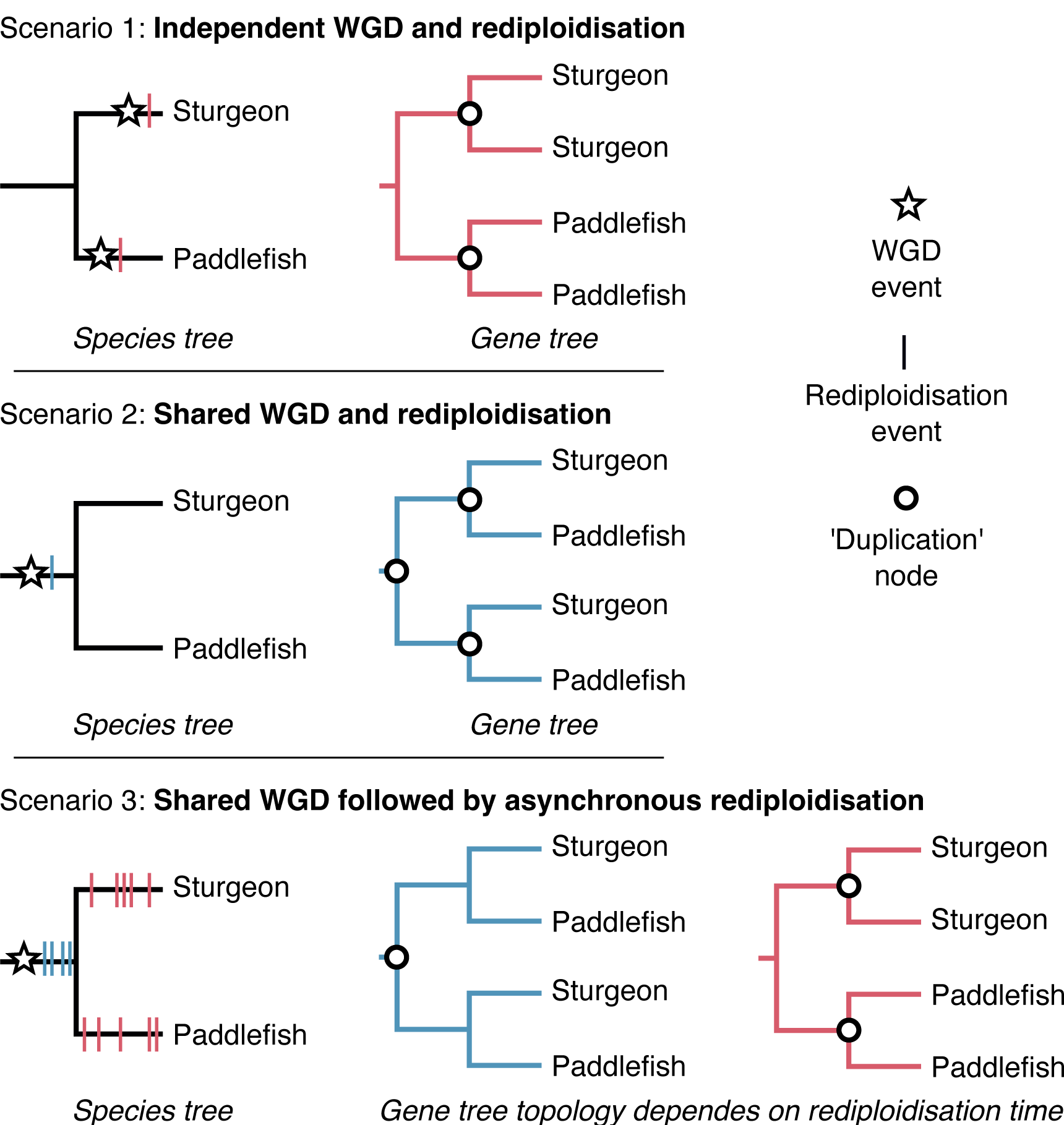
Scenarios of WGD and rediploidisation timing relative to the Sturgeon-Paddlefish divergence and their expected topologies for ohnolog-pair gene trees. Scenario 1 is the widely accepted hypothesis of independent WGD events in the sturgeon and paddlefish lineages. Scenario 2 is a shared ancestral WGD with complete rediploidisation prior to lineage divergence. Scenario 3 extends Scenario 2 by considering the possibility of speciation happening during a prolonged, asynchronous rediploidisation process following a shared WGD event. In this case genes rediploidising prior to speciation will follow the gene tree expected under scenario 2 while those rediploidising after speciation (i.e. lineage-specific rediploidisation) will follow the gene tree expected under Scenario 1. This is distinguishable from independent small-scale duplication using the expectation that ohnolog pairs largely retain ancestral collinearity between non-overlapping duplicate chromosomal regions.

Here, adopting a phylogenomic approach, we reconsider the timing of WGD(s) relative to the sturgeon-paddlefish divergence, accounting for the possibility of lineage-specific rediploidisation after a shared WGD. Taking care to distinguish our results from phylogenetic error, and drawing on conserved synteny, we provide strong evidence for a single ancestral autopolyploidy close to the Permian-Triassic extinction event, followed by extensive lineage-specific rediploidisation, resulting from the ancestral acipenseriform genome remaining predominantly tetraploid at the time of speciation.

## Results

### Recovery of ohnolog pair subsets diverging before and after speciation

Past studies assessing the timing of WGD(s) in acipenseriform history have sought a single consensus ohnolog divergence time relative to speciation^8,35,44^. This approach considers two scenarios as plausible: (1) if a plurality of ohnolog gene trees recover independent duplication nodes after the sturgeon-paddlefish divergence then sturgeons and paddlefish are assumed to have undergone independent WGDs (currently accepted hypothesis) (**Fig. 1**, Scenario 1); and (2) if a plurality of ohnolog gene trees show duplication nodes predating the sturgeon-paddlefish divergence then a single ancestral WGD event can be assumed (typically rejected hypothesis) (**Fig. 1**, Scenario 2). These interpretations implicitly assume all ohnologs share the same rediploidisation timing. Here, we consider a third plausible scenario, as previously observed in salmonid genomes^7,21^: (3) a shared WGD followed by a prolonged rediploidisation process that starts before but continues after speciation. This predicts the presence of two distinct subsets of ohnolog gene trees, one with duplication nodes prior to the sturgeon-paddlefish divergence and the other with independent duplication nodes after speciation (**Fig. 1**, Scenario 3).

To distinguish between these scenarios we added to a set of high confidence ohnologs previously identified in the sturgeon genome by integrating phylogenetic and syntenic evidence^7^. Specifically, we incorporated a broad sampling of proteomes from jawed vertebrate genomes, including from newly available non-teleost ray-finned fish. This allowed us to define 5,439 gene families containing high confidence ohnolog pairs in both sturgeon and paddlefish. Analysing maximum likelihood gene trees for each family we found that the gene tree harbouring independent duplication nodes was the most common topology (hereafter: ‘PostSpec’, for Post-Speciation duplication node, as in Scenario 1 and Scenario 3-right), being recovered 2,074 times (38.13% of all trees; **Fig. 2**). The alternative ohnolog pair topology with a shared duplication node (‘PreSpec’ for Pre-Speciation, as in Scenario 2 and Scenario 3-middle) was recovered 1,448 times (26.62% of all trees; **Fig. 2**). The remaining gene trees (1,917, 35.25%; ‘Other’ for topologies other than PostSpec or PreSpec) failed to recover either of these topologies.

**Figure 2.**
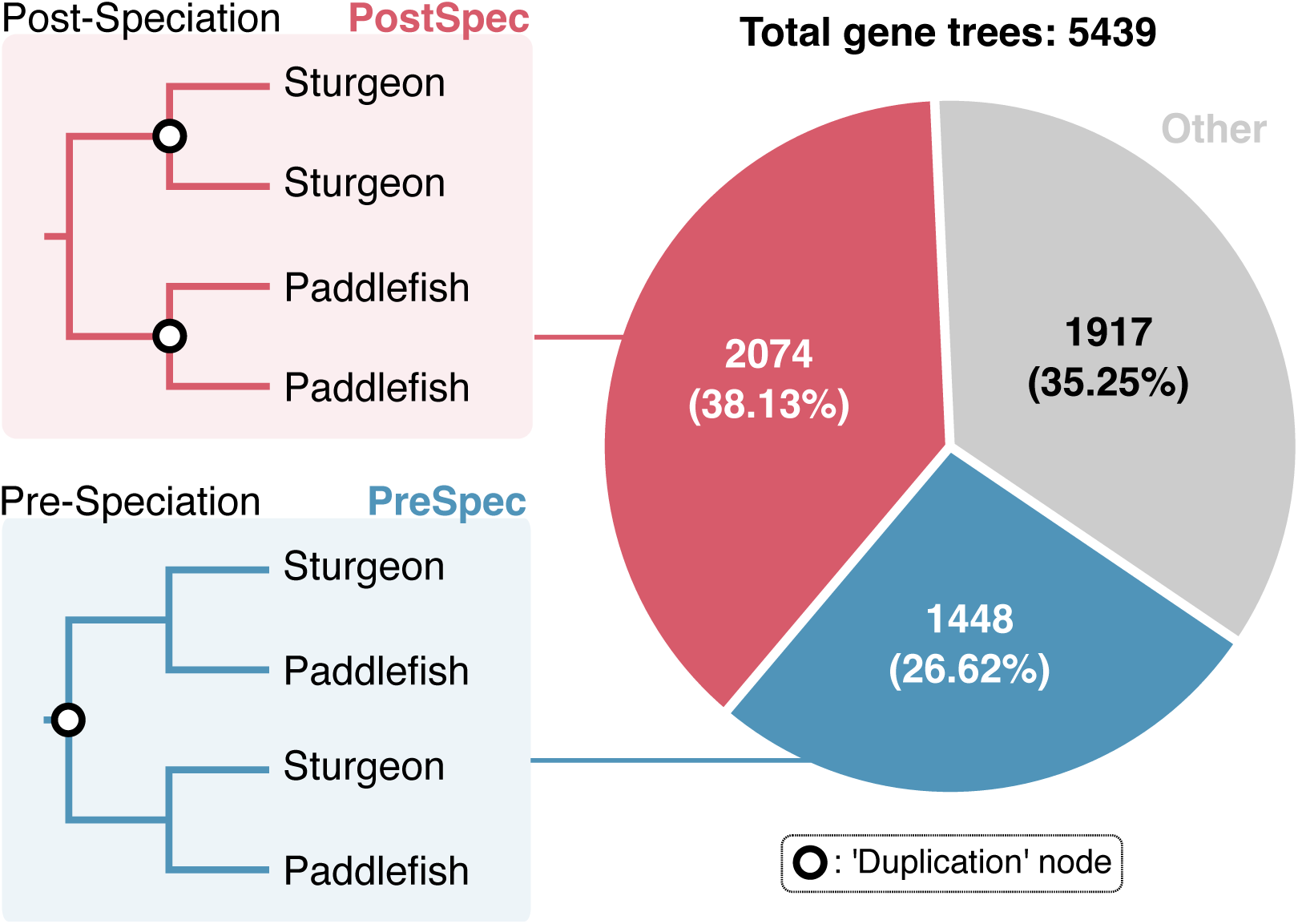
Recovery of key sturgeon-paddlefish ohnolog pair topologies.

The frequent recovery of the PostSpec topology likely explains why previous studies inferred that sturgeons and paddlefish underwent independent WGDs^8,35,38,44,45^ (Scenario 1; **Fig. 1**), however the high prevalence of the PreSpec topology and ‘Other’ topologies in our analyses requires explanation. Firstly, the frequent recovery of both PreSpec and PostSpec topologies is consistent with a shared WGD followed by prolonged rediploidisation extending past the sturgeon-paddlefish speciation (Scenario 3; Fig. 1). In this case, the PreSpec and PostSpec topologies map directly to the ancestral and lineage-specific ohnolog resolution models (dubbed ‘AORe’ and ‘LORe’) previously described in salmonids^24^. However, given the high proportion of inferred ‘Other’ topologies it is important to consider whether the variability in ohnolog divergence time estimates is impacted by phylogenetic error. Similarly, notwithstanding our efforts to define a set of high confidence ohnologs, it is also important to ensure that small-scale duplication events do not drive recovery of either the PreSpec or PostSpec topologies.

### Variation in ohnolog divergence time is not a product of phylogenetic error

To determine the possible impact of phylogenetic error on our findings, we considered factors that may have influenced the initial tree topologies recovered. The critical branching pattern informing our competing hypotheses is the clade in each rooted gene family tree comprising the paddlefish and sturgeon ohnolog pairs (i.e. a 4 gene subtree). First, we considered the recovery of three broad topology categories (PostSpec, PreSpec, ‘Other’; **Fig. 2**) in light of the 15 possible rooted topologies that a four-taxon tree can take. One of these 15 topologies maps to PostSpec, two to PreSpec, and the remaining 12 to ‘Other’ topologies (**Fig. 3A**). However, these ‘Other’ topologies naturally fit into two categories; ‘PostSpec-like’, and ‘PreSpec-like’, each of which were recovered at a frequency in line with their closest main topology (i.e. PostSpec/PreSpec), and require only a single branch change (which could be explained by a minor inference error) to be recovered as PostSpec or PreSpec, respectively (Fig. 3A). Further, such minor topology differences would be indistinguishable from their closest main topology (PostSpec/PreSpec) if the trees were unrooted (**Fig. 3A, Fig. 3B**). The PostSpec and PreSpec topologies, but not ‘Other’ topologies are recovered more frequently than would be expected by random chance (i.e. assuming a 1 in 15 chance for any given topology; **Fig. 3A**). This indicates a strong signal for PreSpec and/or PostSpec but not for ‘Other’ topologies.

**Figure 3.**
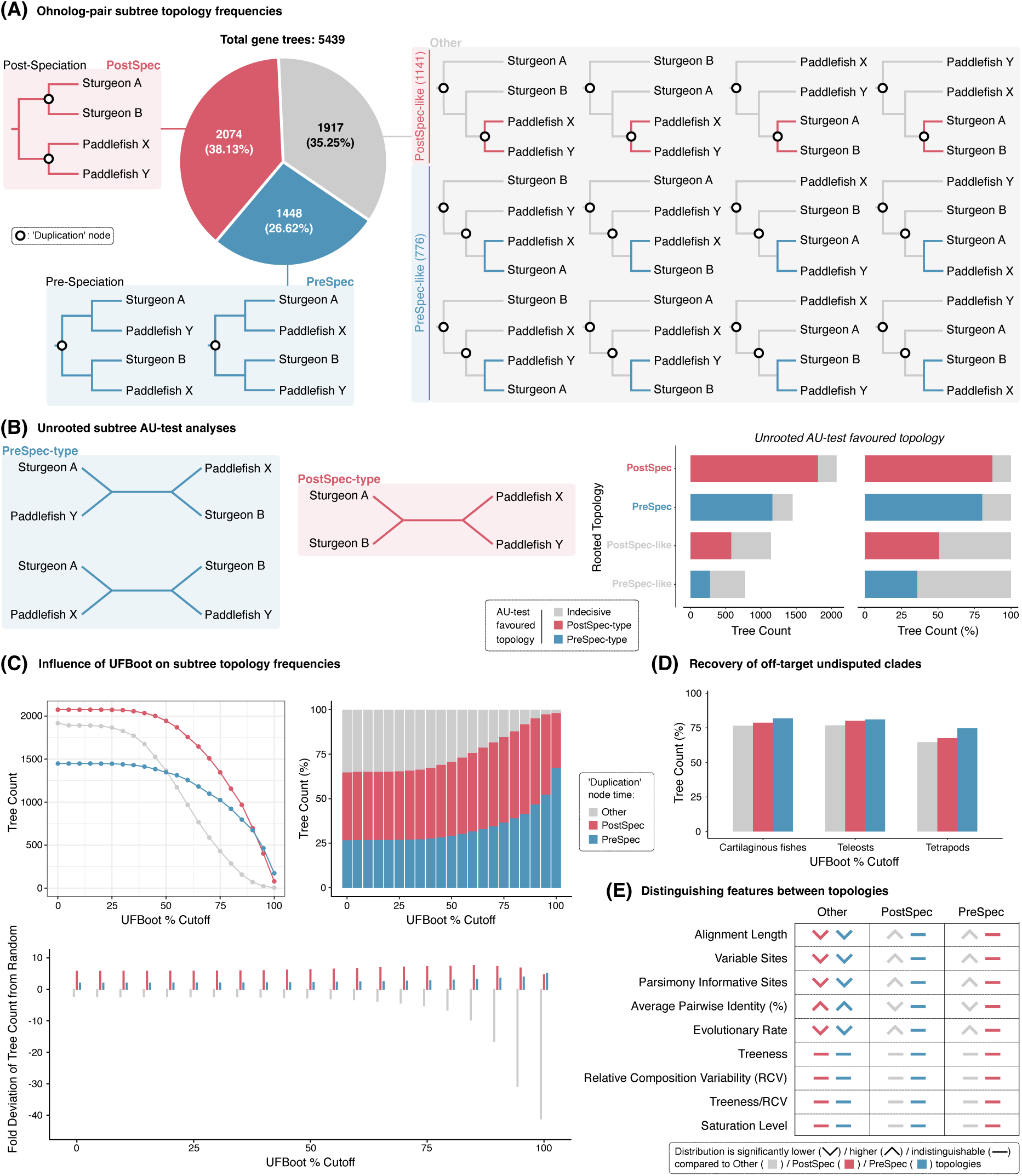
Ohnolog pair gene tree topologies and investigation of possible sources of phylogenetic error. (A) Categorisation of the 15 possible rooted sturgeon-paddlefish subtrees with ‘duplication’ nodes coming before (‘PreSpec’) or after (‘PostSpec’) these species diverged, and ‘Other’ trees that only partially match one of these scenarios (either, ‘PreSpec-like’, or ‘PostSpec-like’). The pie chart quantifies the relative frequency at which each topology was recovered. (B) Three possible unrooted sturgeon-paddlefish subtrees (two ‘PreSpec-type’, left; and one ‘PostSpec-type’, centre), and Approximately Unbiased (AU)-test (right) of tree reliability to determine how frequently datasets from each category of rooted subtree described in part (A) can decisively reject a given unrooted topology category ‘type’ and thereby favour the other. (C) Rooted subtree topology category (broken down at the ‘Other’, ‘PostSpec, ‘PreSpec’ level) count (top left), percentage (top right), and fold deviation of the tree count per category from random expectations (i.e. as estimated if each of the 15 rooted trees were recovered equally frequently; bottom) under increasingly strict UFBoot percentage cut-offs, such that both UFBoot percentages in a given subtree must be greater than or equal to the cut-off for that tree to be retained. (D) Percentage of trees fitting each sturgeon-paddlefish subtree category that recover other key undisputed clades. (E) Summary of significant differences across sequence alignment, modelling, and tree-based statistics for each subtree category (Figure S1 provides violin/box plots with p-values).

To confirm the strength of signal supporting each topology we performed unrooted Approximately Unbiased (AU)-tests^47^ on the sturgeon-paddlefish ohnolog pair subtrees, considering the three possible unrooted topologies of a four taxon tree; one PostSpec-type (the unrooted equivalent of both PostSpec and PostSpec-like), and two PreSpec-type (the unrooted equivalent of both PreSpec and PreSpec-like) (**Fig. 3B**). The results indicate that ohnolog pairs recovering the rooted PostSpec and PreSpec topologies are more robust. Specifically, they frequently reject the unrooted alternative topology type; whereas those that recovered the rooted PostSpec-like and PreSpec-like ‘Other’ topologies reject the unrooted alternative topology type less frequently (**Fig. 3B**). Although the PreSpec/PostSpec-like ‘Other’ datasets are more indecisive, the matching unrooted topology type is almost never rejected in favour of the unrooted alternative topology across any of the four rooted topology sets (**Fig. 3B**). Rather than one or other topology providing a consensus, these results are consistent with significant support for both the PostSpec and PreSpec topologies within our wider ohnolog pair dataset, and with the ‘Other’ topology, being derived from less informative gene family alignments.

As an additional test of tree robustness we assessed the impact of filtering trees based on increasingly stringent branch support cut-offs (i.e. Ultrafast Bootstrap [UFBoot]^48^) within the sturgeon-paddlefish ohnolog pair subtree. As stringency increases, we observe a ‘drop out’ of all tree topologies (**Fig. 3C**). However, this is most severe for ‘Other’ topologies, which are rarely recovered at high stringency, being recovered >40 times less often than random at the strictest cut-off (UFBoot=100%) (**Fig. 3C**). On the other hand, PostSpec and PreSpec topologies were recovered much more often than expected at random, regardless of UFBoot cut-off, with the PreSpec topology overtaking PostSpec as the most frequently recovered topology at UFBoot >=95% (**Fig. 3C**). This further confirms a strong, non-random, signal for both the PostSpec and PreSpec topologies in our full ohnolog pair gene family dataset.

Next, as a proxy for whether we can expect the sturgeon-paddlefish subclade to be recovered accurately, we compared the ability of gene trees supporting each sturgeon-paddlefish ohnolog pair topology to recover other well-accepted clades^49,50^, i.e. cartilaginous fishes, tetrapods, teleosts (**Fig. 3D**). If a gene tree fails to recover known, well-supported clades, it may be indicative of generally low phylogenetic signal. Although no major differences were observed between the three topology categories, PreSpec trees consistently performed best, and ‘Other’ the worst (**Fig. 3D**).

Having confirmed the robustness of the phylogenetic signal, we sought to test whether systematic errors might drive a consistent and misleadingly strong signal for either of PostSpec and PreSpec topologies. We analysed a variety of statistics^51,52^ at sequence alignment, modelling and tree topology levels (**Fig. 3E, Fig. S1**). ‘Other’ tree topologies derive from shorter multiple sequence alignments with fewer substitutions per site (i.e. higher average pairwise identity, slower evolutionary rate^53^, and fewer variable and parsimony informative sites) than PostSpec or PreSpec topologies (**Fig. 3E, Fig. S1**). The combination of these factors presumably limits phylogenetic signal, consistent with the idea that ‘Other’ topologies result from weakly-supported, minor phylogenetic errors. We observed no significant differences across any statistics considered between the PreSpec and PostSpec topologies (**Fig. 3E, Fig. S1**). Importantly, although they have more substitutions per site than ‘Other’ trees, PostSpec and PreSpec datasets do not show signs of being more susceptible to systematic errors, having a comparable balance of substitutions on internal and external tree branches (treeness), and similar compositional variability and substitutional saturation levels to ‘Other’ datasets (**Fig. 3E, Fig. S1**)^52,54,55^.

Lastly, site-heterogeneous models can help to alleviate systematic error-induced branching artefacts in phylogenetic analysis^56^. Testing their use on all gene families that had maximal support values (UFBoot=100%) within the sturgeon-paddlefish ohnolog pair subclade never resulted in a topology change, while support values never dropped below UFBoot=97% (**Fig. S2**).

Together these analyses indicate that neither the PostSpec and PreSpec topologies derive from error, indicating strong and reliable support for ohnologs diverging both before and after the sturgeon-paddlefish divergence, while suggesting that ‘Other’ tree topologies are a product of comparatively limited phylogenetic signal.

### Conserved synteny supports a shared WGD followed by prolonged and asynchronous rediploidisation

The autopolyploid rediploidisation process is thought to involve numerous physically independent genomic rearrangement events across the genome^24^. Assuming that these rearrangements are not restricted to single genes^24,26^, and that subsequent rearrangements are not extensive, large blocks of neighbouring genes sharing common rediploidisation histories should be visible as largely non-overlapping syntenic blocks on different chromosomes, and present in both lineages. This is not unlike the history of suppression of recombination during the evolution of mammalian sex chromosomes, where genome rearrangements are associated with the onset of locus divergence on the X and Y and resulted in contiguous ‘strata’ of genes that share an X-Y divergence time^14,57^. If our phylogenetic results arise from a shared WGD followed by a prolonged rediploidisation spanning both the shared and lineage-specific branches, such divergence-time-stratified synteny blocks should be highly evident, especially considering that acipenseriform genomes evolve slowly and show limited reorganisation after WGD^7,8,35^.

Plotting ohnolog pairs within and across the sturgeon and paddlefish genomes revealed that ohnologs from both the PreSpec and PostSpec categories (**Fig. 2, 3**) are not randomly distributed along the genome. Instead ohnolog pairs with shared divergence dates relative to speciation (PreSpec or PostSpec) are found in syntenic blocks along large uninterrupted sections of chromosomes (and possibly even entire small chromosomes) (**Fig. 4**). For example, long PreSpec synteny blocks are conserved across both genomes on the six largest chromosomes (which form three WGD-derived pairs across both species^7,8^; **Fig 4C**), making small-scale segmental duplication prior to lineage-specific WGD an implausible explanation for these topologies, and adding further support to the hypothesis that they reflect true evolutionary signal stemming from WGD. In all, these data are parsimoniously explained by a single ancestral WGD followed by extensive ancestral and lineage-specific rediploidisation in sturgeon and paddlefish evolution.

**Figure 4.**
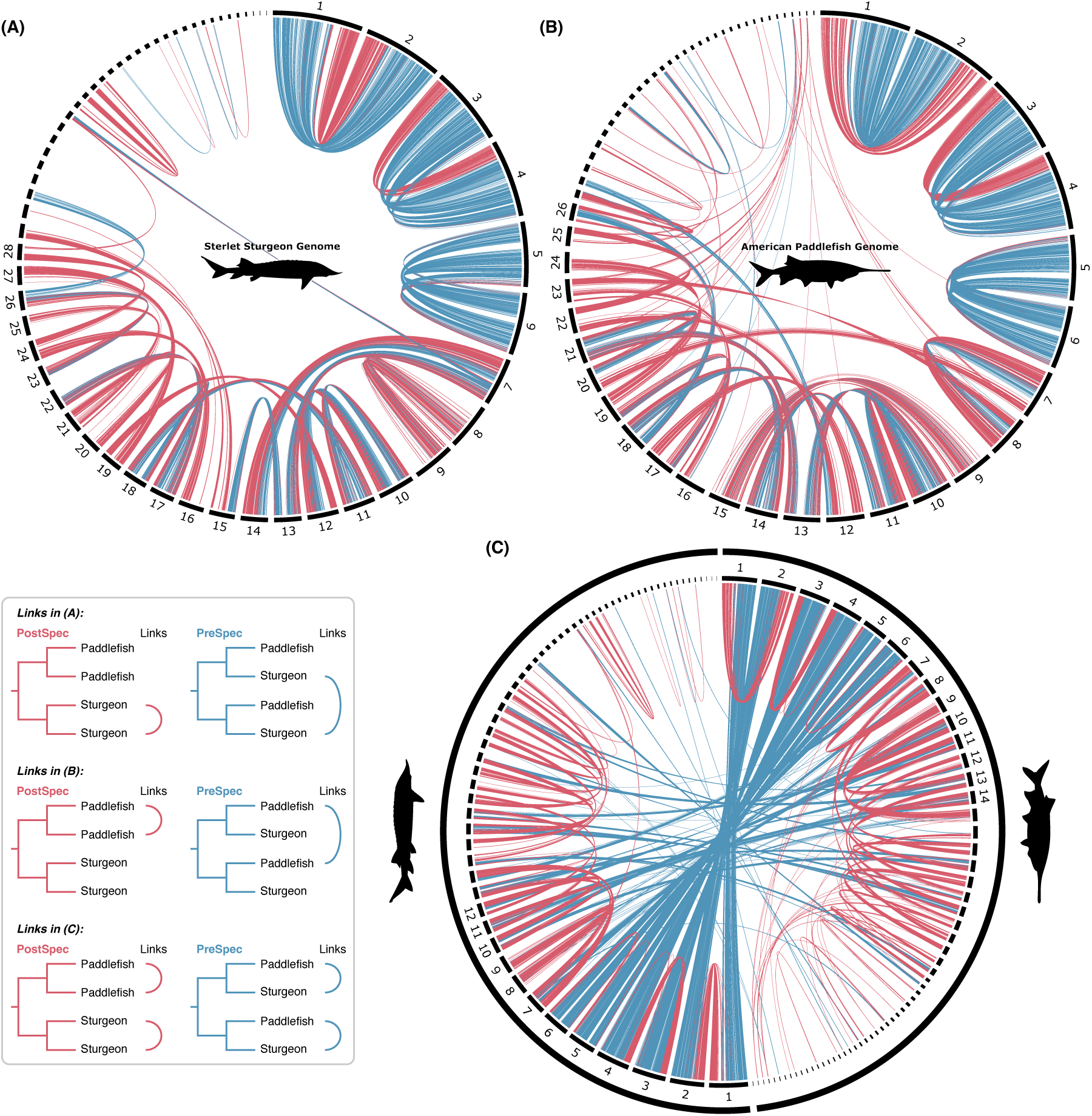
Synteny patterns of ‘PreSpec’ and ‘PostSpec’ ohnolog pairs in the paddlefish and sturgeon genomes. Circos plots of the sterlet sturgeon genome (A) and the American paddlefish genome (B) showing the chromosomal locations of ohnolog pairs, with links coloured according to the PreSpec (blue) or PostSpec (red) tree topology. Microchromosomes <20Mb are not labelled. (C) Circos plot of ohnolog-pairs in both the sturgeon and paddlefish genomes, with intra-specific PostSpec links (red) and inter-specific PreSpec links (blue). Only macrochromosomes >40Mb from each species are labelled.

This result stands even when examining only the macrochromosomes (>40Mb) (**Fig. S3**) and for ohnolog trees with progressively stricter statistical support (i.e. UFBoot cut-off scores of ≥50%, ≥75%, and 100%) (**Fig. S4**). Meanwhile, ohnolog trees from the ‘Other’ category tend to occupy genomic regions harbouring genes with the most similar topologies (i.e. PreSpec-like alongside PreSpec, and PostSpec-like alongside PostSpec; **Fig. S5**) as expected if these topologies primarily arise from minor errors.

### A Permian-Triassic lower bound for the sturgeon-paddlefish WGD

Asynchronous rediploidisation temporally separates ohnolog divergence from WGD, obscuring the dating of autopolyploidy events^24,26,58^. Although imperfect, the most reliable lower bound estimate for the ancestral sturgeon-paddlefish WGD event can be estimated from ohnolog pairs that rediploidised prior to the sturgeon-paddlefish divergence, as these will have diverged closer in time to the WGD event ^58^. With this in mind, to estimate a lower bound timing for the WGD we took a Bayesian phylogenomic approach^59^ using concatenated ohnolog pairs^32,58^ based on the set of 81 gene trees that maximally supported shared WGD and did not include duplicates in other species. We analysed five distinct datasets, always including all 81 gene families but randomly shuffling ohnologs from a pair for arbitrary assignment as the ‘A’ or ‘B’ copy for concatenation, to avoid bias and assess robustness of results to alternative concatenations^58^. Analysing these datasets with an autocorrelated relaxed molecular clock^60^, and using the site-heterogeneous CAT-GTR substitution model^61^, we infer a divergence time of ∼171.6 Ma (average mean of all five random concatenations; 95% credibility interval range: ∼124.1-203.3 Ma) for the split of sturgeons and paddlefish (i.e. crown Acipenseriformes; considering both ohnolog pairs) (**Fig. 5, Fig. S6**). Following the splitting of Chondrostei (of which sturgeons and paddlefish are the only living representatives) and Neopterygii ∼367.8 Ma (average mean of all five random concatenations; 95% credibility interval range: ∼360.6-374.8 Ma), we estimate a lower bound for the shared sturgeon-paddlefish WGD at ∼254.7 Ma (average mean of all five random concatenations; 95% credibility interval range: ∼207.1-289 Ma). (**Fig. 5, Fig. S6**). Thus, the mean Bayesian estimate for the timing of the ancestral sturgeon-paddlefish WGD lower bound sits close to the Permian-Triassic (P-Tr) boundary mass extinction event ∼251.9 Ma.

## Discussion

Previous studies have favoured independent WGD events in the sturgeon and paddlefish lineages, despite their close phylogenetic relationship^8,35,38,44,45^. By accounting for the possibility of a single autopolyploidy event followed by lineage-specific rediploidisation^24^, our results reject independent WGDs, revealing that an ancestral WGD was followed by speciation at a time when ∼50-66% of the genome remained tetraploid. This high proportion of tetraploidy at the time of speciation provides an explanation for past studies incorrectly inferring independent WGD events^8,35,38,44,45^ and has implications for acipenseriform evolution and biology.

**Figure 5.**
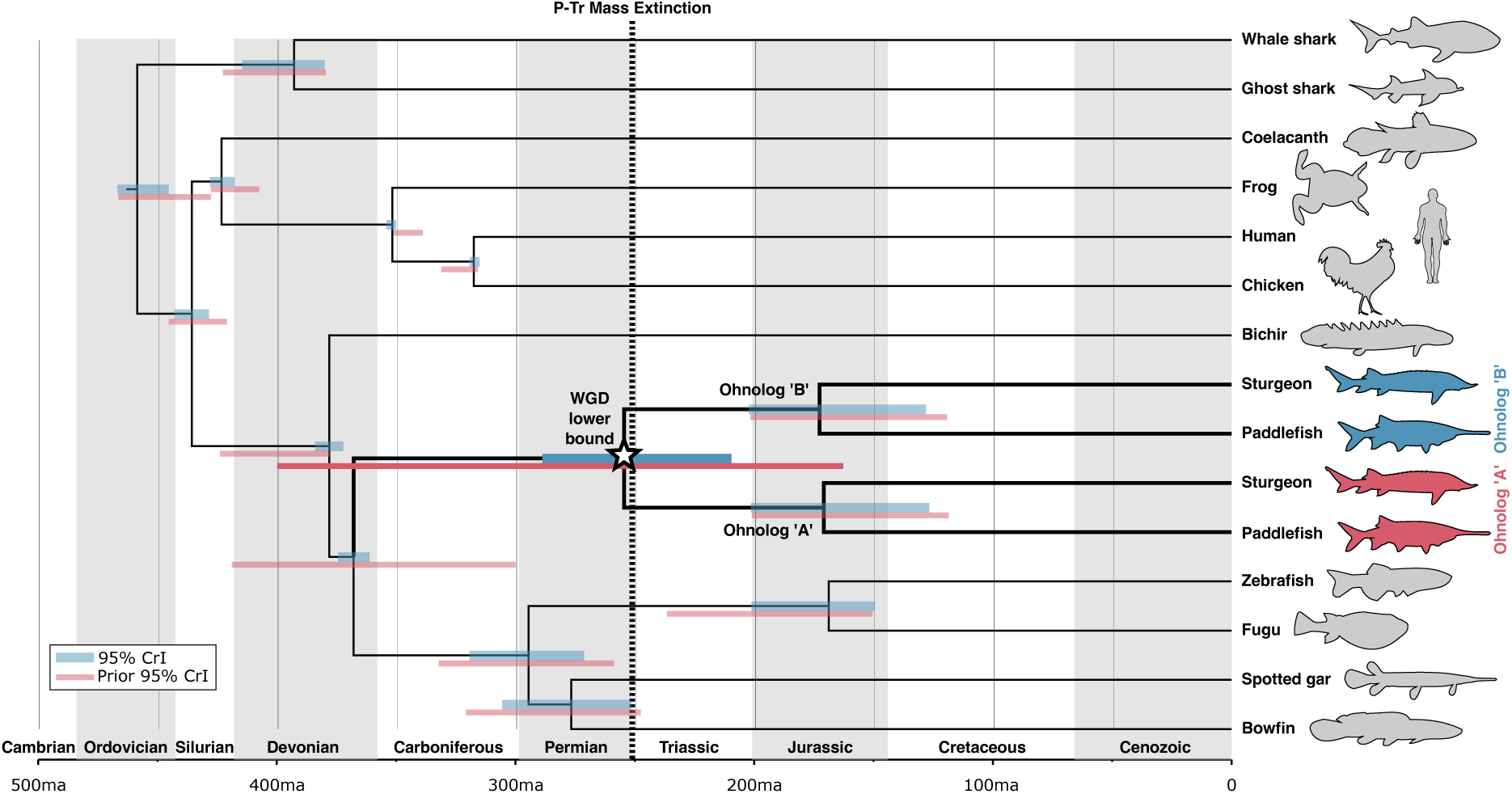
Phylogenomic dating of the shared sturgeon-paddlefish WGD lower bound. Ohnolog copies were randomly classified as ‘A’ or ‘B’ and concatenated for phylogenomic analysis. The jawed vertebrate Bayesian phylogenomic timetree from one of five random concatenations is shown (for all five see Fig. S6). The 95% CrI (credibility interval) is shown for each node in blue. The 95% CrI results from an independent analysis under the prior are shown below each node in red, verifying that our priors on divergence times are sufficiently diffuse to have avoided restricting our results to the inferred WGD lower bound timing in the main analyses.

Though not framed in the context of a shared WGD, similarity in the evolution of the sturgeon and paddlefish genomes has been previously noted^8^. For example, the six largest chromosomes (three WGD-derived chromosome pairs) appear to show particularly strong chromosomal homology between sturgeon and paddlefish (**Fig. 4C**)^8^; and we find that these same chromosomes have primarily undergone rediploidisation before the sturgeon-paddlefish divergence. For these and other regions of the genome that have undergone rediploidisation before speciation, the onset of ohnolog sequence and functional divergence (beyond allelic variation), will also have been ancestral to both extant acipenseriform lineages. This partially shared rediploidisation history after shared WGD likely explains at least some of the proposed similarity in genome evolution between paddlefish and sturgeons and perhaps contributes to their ability to hybridize^8,62^.

Conversely, other parts of the genome—those chromosomes and regions of chromosomes that were tetraploid when sturgeons and paddlefish diverged— rediploidised independently in each lineage, with different (and differently sized) regions, and hence different sets of genes, rediploidising at different times. Those ohnologs that resolved from alleles after the speciation must have also undergone any sub-/neo-functionalisation/regulation independently^24^. Given that over half of the genome appears to have rediploidised after the sturgeon-paddlefish divergence, it is likely that these independently rediploidised ohnologs, and the networks they form, contribute substantially to the unique biology of each lineage. For example, a recent study of the oxytocin and vasotocin receptor (OTR/VTR) gene family, which play a variety of roles, including in social behaviour and reproduction, found that these duplicated genes emerged consistent with independent WGDs in each lineage^63^. Our results instead indicate that the OTR/VTR genes might be better interpreted as following the LORe model^24^, having rediploidised independently in sturgeons and paddlefish.

Our estimate that ∼50-66% of the duplicated genome remained tetraploid at the point of the sturgeon-paddlefish divergence i.e. ∼80 million years post-WGD, is much more drawn-out than equivalent estimates for salmonids (∼60-70% rediploidisation completed after ∼50 million years^24,26^) and teleosts (rediploidisation largely resolved after ∼60 million years^13,64^) (**Fig. 5, Fig. S6**)). We suggest that the apparently slower evolutionary rate (in terms of both substitutions and rearrangements^7,8^) in sturgeons and paddlefish contributed to a more prolonged rediploidisation period in Acipenseriformes compared to faster evolving teleosts. Despite this slower rate of genome rearrangement and rediploidisation, and the presence of large blocks of genes sharing consistent rediploidisation history in our analyses, we find ohnolog blocks that diverged before and after speciation on the same chromosomes, similar to observations in salmonids^24,26^. This appears to have often occurred through temporally isolated intrachromosomal rearrangement events, perhaps facilitating suppression of homologous recombination and allowing resolution of ohnologs from alleles for that genomic segment. This scenario is akin to the stepwise formation of evolutionary ‘strata’ on mammalian sex chromosomes^14,57^, and adds a layer of complexity atop the existing model of segmental rediploidisation proposed for sturgeon^7^.

Rediploidisation occurring asynchronously means that common approaches (e.g. phylogenomics or molecular clock-based analyses) used to directly estimate the absolute date of autopolyploid WGD events are unavoidably problematic, as they conflate ohnolog rediploidisation time(s) with the WGD itself^24,26,58^. Our analyses indicate that the sturgeon-paddlefish WGD must predate the emergence of these lineages by at least long enough for ∼33-50% of the genome to have rediploidised. Our WGD lower-bound timing (∼254.7 Ma) is older than previous estimates^7,8,38,44,45^ and suggests the WGD occurred very early in acipenseriform evolution, if not before the emergence of the order as a whole within Chondrostei. This timing implies that stem Acipenseriformes (such as <u>†</u>Peipiaosteidae and <u>†</u>Chondrosteidae)^65–68^ likely split from the ancestor of extant Acipenseriformes with genomes still early in the rediploidisation process.

Our results support a potential role for polyploidy in Acipenseriformes surviving the P-Tr mass extinction event. It has been proposed that WGDs in plants may confer tolerance and adaptability to extreme environmental conditions, increasing fitness in the face of mass extinction events^34,69^. Our dating of the ancestral sturgeon-paddlefish WGD is consistent with a model where the flexibility and functional redundancy intrinsic to a genome in the early stages of autopolyploid rediploidization contributed to the survival and success of the Acipenseriformes through the P-Tr mass extinction. The discovery of a mix of ancestral and lineage-specific rediploidisation in both teleost^13,27,28,24,26^ and non-teleost ray-finned fish lineages, suggests it is a general phenomenon after WGD, at least for autopolyploids. Because any individual gene cannot be considered duplicated until recombination is suppressed, such a scenario generates genomes consisting of a mosaic of shared and lineage-specific gene duplications, even though they originated from a single genome duplication. This complex relationship may help to explain the long-standing difficulty in resolving the number and timing of WGDs in early vertebrate evolution^9,11,12,70–72^.

Extensive lineage-specific rediploidisation has major implications for our understanding of genome evolution following polyploidy and for our interpretation of evolution of duplicate genes including their role in adaptive evolution. This new framework for the analysis and interpretation of evolution following WGD will prompt a re-examination of other autopolyploidy events, including the founding WGD at the base of all vertebrates.

## Methods

### Ohnolog-pair datasets

In their analysis of the sterlet sturgeon genome, Du et al. (2020)^7^ defined a high-confidence ohnolog pair dataset for the species, and this forms an initial basis for our analyses. To add paddlefish (GCF_017654505.1)^8^ ohnologs to this dataset we used OrthoFinder^73^ (v 2.5.4) to infer phylogenetic hierarchical orthogroups (PHOGs). For OrthoFinder analyses we also included a set of proteomes from species spanning jawed vertebrate phylogeny, including the ghost shark (*Callorhinchus milii*; GCF_000165045.1)^74^ and whale shark (*Rhincodon typus*; GCF_001642345.1)^75^ from Chondrichthyes, and human (*Homo sapiens*; GCF_000001405.39), domestic chicken (*Gallus gallus*; GCF_000002315.6), African clawed frog (*Xenopus tropicalis*; GCF_000004195.4), and coelacanth (*Latimeria chalumnae*; GCF_000225785.1) from Sarcopterygii. Within Actinopterygii we selected zebrafish (*Danio rerio*; GCF_000002035.6), fugu (*Takifugu rubripes*; GCF_901000725.2), spotted gar (*Lepisosteus oculeatus*; GCF_000242695.1)^39^, and bowfin (*Amia Calva*; JAAWVP01.1)^35^ as representatives of Neopterygii, the sister group to sturgeons and paddlefishes, and grey bichir (*Polypterus senegalus*; GCF_016835505.1)^35^ as their combined sister group. The longest protein sequence for each gene was used for these analyses where alternative transcripts were annotated. Default OrthoFinder settings were used, with two exceptions. First, we specified a species tree, in line with accepted jawed vertebrate relationships^7,8,35,76^, to augment orthology inference: “((GhostShark,WhaleShark),((Coelacanth,(Frog,(Human,Chicken))),(Bichir,((Paddlefish,Sturgeon),((Zebrafish,Fugu),(SpottedGar,Bowfin))))));”. Second, the OrthoFinder ‘-y’ flag was specified to further split PHOGs that underwent duplications after the jawed vertebrate ancestor into separate PHOGs. In post-processing of OrthoFinder results, we then performed an extra check by extracting only PHOGs including sequences from as many species as possible, while always including two sequences each from sturgeon and paddlefish. This was achieved by extracting the set of sequences descended from the most ancient ancestral species node in the OrthoFinder reconciled gene tree to not include additional sturgeon or paddlefish sequences (based on the ‘.tsv’ files in the OrthoFinder ‘Phylogenetic_Hierarchical_Orthogroups’ folder). As a simple example, in an orthogroup with a gene duplication in the ancestor of Actinopterygii, with both paddlefish and sturgeon ohnologs being retained in both actinopterygian duplicates, our approach results in two resultant PHOGs, split at the level of Actinopterygii, with neither including sequences from the other duplicate or their co-orthologs from Sarcopterygii or Chondrichthyes, and both containing two sturgeon and two paddlefish sequences each.

These PHOGs were then filtered to retain only those that matched a previously inferred sturgeon high-confidence ohnolog pair^7^, as well as those including at least one outgroup to allow rooting of the sturgeon-paddlefish ohnolog pair subtree, and hence inference of duplication node time relative to speciation. We also excluded gene families where both ohnologs in either paddlefish or sturgeon were present on the same chromosome, or where any paddlefish or sturgeon sequence was present on a scaffold not assigned to one of the 60 sturgeon or paddlefish chromosomes. Lastly, we checked that the sturgeon and paddlefish sequences formed a monophyletic group in inferred PHOG gene trees (see below section on Ohnolog duplication time inference). 5,439 PHOGs met these criteria (and form our ohnolog-pair set), of which 5,372 also included at least one sequence each from Neopterygii and a more distantly related outgroup, all but ensuring that the two paddlefish and sturgeon sequences diverged after splitting from Neopterygii.

In all, this should have resulted in a dataset heavily enriched for ohnolog pairs in sturgeon and paddlefish. However, we note that the set includes only ohnolog pairs where both ohnologs are retained in both species and excludes PHOGs that have undergone additional duplications or losses in sturgeons, paddlefish, or their ancestral stem lineage. Similarly, it is possible that a very small subset of our ohnolog pairs may derive from complex scenarios of multiple ohnolog losses and inter-chromosomal duplications that have evaded our filters, as well as the doubly conserved synteny evidence for the original sturgeon ohnolog set, and left a pair of ohnolog ‘look-a-likes’ in paddlefish and sturgeon. However, we expect PHOGs where this has occurred to be exceedingly rare, if present at all, in our final 5,439 sturgeon-paddlefish ohnolog-pair dataset.

### Ohnolog duplication time inference

To estimate rediploidisation time (i.e., ‘duplication’ node time) relative to speciation for each sturgeon-paddlefish ohnolog-pair we performed phylogenetic analyses for each gene family from the pre-monophyly filtered sturgeon-paddlefish ohnolog pair PHOGs described above (5,590 trees). Multiple sequence alignments were performed with MAFFT v7.487 with the ‘--auto’ flag specified, and the ‘--anysymbol’ flag for those datasets that included sequences containing selenocysteine (symbol ‘U’). Phylogenetic inference was performed using IQ-tree^77^ (v. 2.1.4-beta COVID-edition), with the ‘-m JTT+G’ flag to use the JTT^78^ amino acid substitution model with four discrete gamma categories, as well as the ‘-bb 1000’ flag to specify 1000 ultrafast bootstrap replicates^48^. The resulting maximum likelihood trees were extracted for pre-processing and duplication time inference. To pre-process these trees, we used the ETE (v3) toolkit^79^ python library to check that sturgeon and paddlefish sequences formed a ‘monophyletic’ clan^80^ in each PHOG gene tree, and then rooted each tree with the most distantly related sequence relative to sturgeons and paddlefish (i.e. typically ghost shark/whale shark). In a separate python script, the ETE toolkit was then used to perform strict gene tree-species tree reconciliation^79,81^ to infer speciation and duplication nodes/events, before classifying (PreSpec, PostSpec, ‘Other’ [PreSpec-like, PostSpec-like]) and summarising the different sturgeon-paddlefish subtree gene tree topologies and frequencies recovered.

### Testing robustness of ohnolog gene tree topologies to error

To interrogate the possibility that either of the PreSpec or PostSpec gene trees derive from error, as well as to better understand the source of the ‘Other’ topologies we performed a suite of analyses. First, we performed unrooted AU-test^47^ analyses in IQ-tree^77^. To do this we classified the three possible unrooted topologies of our four-tipped sturgeon-paddlefish subtree as either PostSpec-type or PreSpec-type based on whether they would become PostSpec(-like) or PreSpec(-like) if rooted. For each of the four rooted topology categories (i.e., PostSpec, PreSpec, and the two ‘Other’ categories of PreSpec-like and PostSpec-like) we extracted the subalignment for the four sturgeon/paddlefish sequences from each PHOG in that category, and then performed an AU-test analysis for each ohnolog-pair set specifying the three possible unrooted topologies. The frequency at which an unrooted topology type was favoured by the AU-test (i.e., the AU-test reject the alternative unrooted topology/topologies) for each rooted tree topology category was then calculated and plotted.

Next, using a python script and the ETE toolkit^79^ we assessed the influence that filtering our ohnolog-pair PHOG counts by increasingly higher ultrafast bootstrap percentage cut-offs (considered for the two support values within the four-tipped paddlefish-sturgeon clade only) would have on rooted topology frequencies. Starting at a cut-off of 0% and incrementing by 5%, up to 100%, we assessed the total counts and percentages of PostSpec, PreSpec, and ‘Other’ tree topologies recovered at each cut-off and then assessed for trends in topology frequency as the cut-off became more stringent. We also calculated and plotted the fold deviation from random (i.e. if all 15 topologies were recovered equally frequently) at which each rooted tree category was recovered for PostSpec (randomly expected 1/15 times), PreSpec (randomly expected 2/15 times) and ‘Other’ (randomly expected 12/15 times) topologies across the same series of ultrafast bootstrap cut-offs and evaluated the trends observed.

To assess whether gene families supporting any topology perform poorly at recovering other generally-accepted clades^49,50^, and hence may be more likely to be misleading, we used the custom ETE toolkit^79^ python script to assess for the monophyly of three widely-accepted clades; Tetrapoda (tetrapods; monophyly of human, chicken, and frog), Teleostei (teleost fishes; monophyly of fugu and zebrafish), and Chondrichthyes (cartilaginous fishes; monophyly of ghost/elephant shark and whale shark). This script also required that at least one sequence was present for each species in that monophyletic clade, meaning some negatives may also derive from gene loss or absence from our inferred PHOG.

To detect signs of possible systematic errors, we compared a range of statistics at the sequence alignment, modelling, and inferred phylogenetic tree levels, between gene trees fitting the ‘PostSpec’, ‘PreSpec’, and ‘Other’ tree topologies using the Wilcox-test with Bonferroni correction in R. Alignment length and average pairwise percentage identity were calculated using the esl-alistat program from the Hmmer package [version 3.1b2; http://hmmer.org]^82^, while the number of parsimony informative sites^52^ was extracted from IQ-tree output. PhyKIT^51^ was used to compute the number of variable sites^52^, the evolutionary rate (i.e. total tree length/number of leaf nodes)^53^, treeness (i.e. sum of internal branch lengths/total tree length)^54^, relative compositional variability (RCV)^54^, treeness/RCV^52,54^, and saturation level^55^.

Lastly, to further explore whether our results could derive from systematic error, we tested the use of precomputed site-heterogeneous mixture models on the set of PHOGs that had maximal support for any sturgeon-paddlefish subclade topology (ultrafast bootstrap = 100% for both support values in the sturgeon-paddlefish subtree). Specifically we tested the fit and influence the UL3^83^, EX_EHO^84^, and JTT+C20^85^ models, which can help to alleviate systematic biases, and often fit single gene family alignments well^86,87^.

### Synteny analysis

The genomic coordinates of each gene in an ohnolog pair were used to anchor links between ohnologs on circos plots (drawn with circos-0.69-9^88^) of the sturgeon and paddlefish genomes. All members of an ohnolog pair were required to be present on the largest 60 chromosomes to be included. For plotting of both species in a single circos plot, PreSpec ohnolog pairs were split into separate ohnologs to be plotted as orthologs between species, while PostSpec ohnolog pairs were plotted as ohnologs within species as per the individual species plots.

### Phylogenomic divergence dating

To estimate a lower bound for the sturgeon-paddlefish WGD, we extracted the set of gene trees recovering the PreSpec sturgeon-paddlefish ohnolog pair topology with maximal support (ultrafast bootstrap = 100% for both support values in the sturgeon-paddlefish subtree). To simplify preparation for phylogenomic analysis and reduce computation time of dating analyses, we then filtered for gene families that were otherwise single copy, resulting in a set of 81 gene families. We generated five distinct datasets by randomly assigning ohnologs from a pair as the ‘A’ or ‘B’ copy prior to concatenation of all 81 existing gene family multiple sequence alignments. This avoids bias from a single arbitrary concatenation, while also permitting assessment of how robust results are to variations in ohnolog concatenations^58^.

Upon concatenation each super-matrix was then filtered using trimAl^89^ (‘-nogaps’) and BMGE^90^ (‘-m BLOSUM62’) to trim gap-rich and saturated sites, after which 42,126 amino acid alignment sites remained in each dataset.

Phylogenomic divergence dating was then performed (on each of the five alternative super-matrices) in Phylobayes^59^ (version 4.1c), specifying the site-heterogeneous CAT-GTR+G4^61^ substitution model along with an autocorrelated lognormal relaxed clock model^60^ (which fits and performs well in jawed vertebrate phylogenomics^76^), and a birth-death prior with soft bounds^91,92^ on fossil calibrations. Fossil calibrations priors (**Table S1**) were set for most nodes across the tree, with the notable exceptions of the sturgeon-paddlefish WGD lower bound timing node, and the node splitting sturgeons and paddlefish (Acipenseriformes) from Neopterygii. Calibrations, including a minimum divergence of 121 Ma^93,94^ for sturgeons and paddlefish, followed Benton et al. (2015)^95^, except for the lower bound on crown Chondrichthyes which was set at 381 Ma^96^. A fixed tree topology (“((GhostShark,WhaleShark),((Coelacanth,(Frog,(Human,Chicken))),(Bichir,(((Paddle fishA,SturgeonA),(PaddlefishB,SturgeonB)),((Zebrafish,Fugu),(Gar,Amia))))));”) was specified based on accepted jawed vertebrate phylogeny^7,8,35,76^ and our inference of a shared WGD, and the Chondrichthyes representatives, ghost shark and whale shark, were set as the outgroup. We verified this topology for each of our five datasets by performing a basic concatenated phylogenomic analysis in IQ-tree^77^ under the JTT+G4 model^78^ with 1000 ultrafast bootstrap replicates^48^.

Each Phylobayes Markov chain Monte Carlo analysis was sampled for at least 10,000 cycles, with the first 5,000 discarded as burn-in before calculation of inferred divergence dates and 95% credibility intervals. Runs under the prior were performed using the same settings, except for swapping to a site-homogeneous Poisson substitution model for computational efficiency since the prior over divergence times is independent of substitution model priors, to verify that the prior on the sturgeon-paddlefish WGD lower bound timing were sufficiently diffuse as to be uninformative.

## Data Availability

All python scripts, ohnolog pairs and associated data, including gene family alignments and trees, error assessment statistics, as well as random concatenation sets and phylogenomic super-matrices are available on figshare.

## Acknowledgements

We thank Dr Matthias Stöck for sharing the sterlet sturgeon genome annotation. AKR is supported by an Irish Research Council Government of Ireland Postdoctoral Fellowship (GOIPD/2021/466). This work was supported by funding from the European Research Council, grant agreement 771419 (AMcL).

## Author contributions

AKR and AMcL devised the study with input from DJM and MKG. AKR carried out analyses. AKR and AMcL analysed and interpreted results. AKR, AMcL and DJM wrote the manuscript.

