## Supplementary Data for "Extensive lineage-specific rediploidisation masks shared whole genome duplication in the sturgeon-paddlefish ancestor"

**File includes Table S1 and Figures S1-S6.**

**Table S1.** Fossil calibrations used in phylogenomic dating of the sturgeon-paddlefish WGD.

| Calibrated node | Minimum Age Calibration (Ma) | Maximum Age Calibration | References |
|---|---|---|---|
| Crown Gnathostomata/Jawed Vertebrates (Human – Whale Shark) | 421 | 465 | [1,2] |
| Crown Osteichthyes (Human - Zebrafish) | 421 | 445 | [1] |
| Crown Sarcopterygii (Human - Coelacanth) | 408 | 428 | [1] |
| Crown Tetrapoda (Human - Frog) | 337 | 351 | [1] |
| Crown Amniota (Human - Chicken) | 318 | 333 | [1] |
| Crown Chondrichthyes (Whale Shark – Ghost Shark) | 381 | 423 | [3] |
| Crown Actinopterygii (Bichir – Zebrafish) | 378 | 423 | [1] |
| Crown Neopterygii (Zebrafish – Spotted Gar) | 250 | 331 | [1] |
| Crown Clupeocephala (Zebrafish - Fugu) | 151 | 235 | [1] |
| Crown Holostei (Spotted Gar - Bowfin) | 250 | 332 | [1] |
| Crown Acipenseriformes (Sturgeon – Paddlefish) [Included twice in analyses for the A and B ohnolog pairs] | 121 | 202 | [1,4,5] |

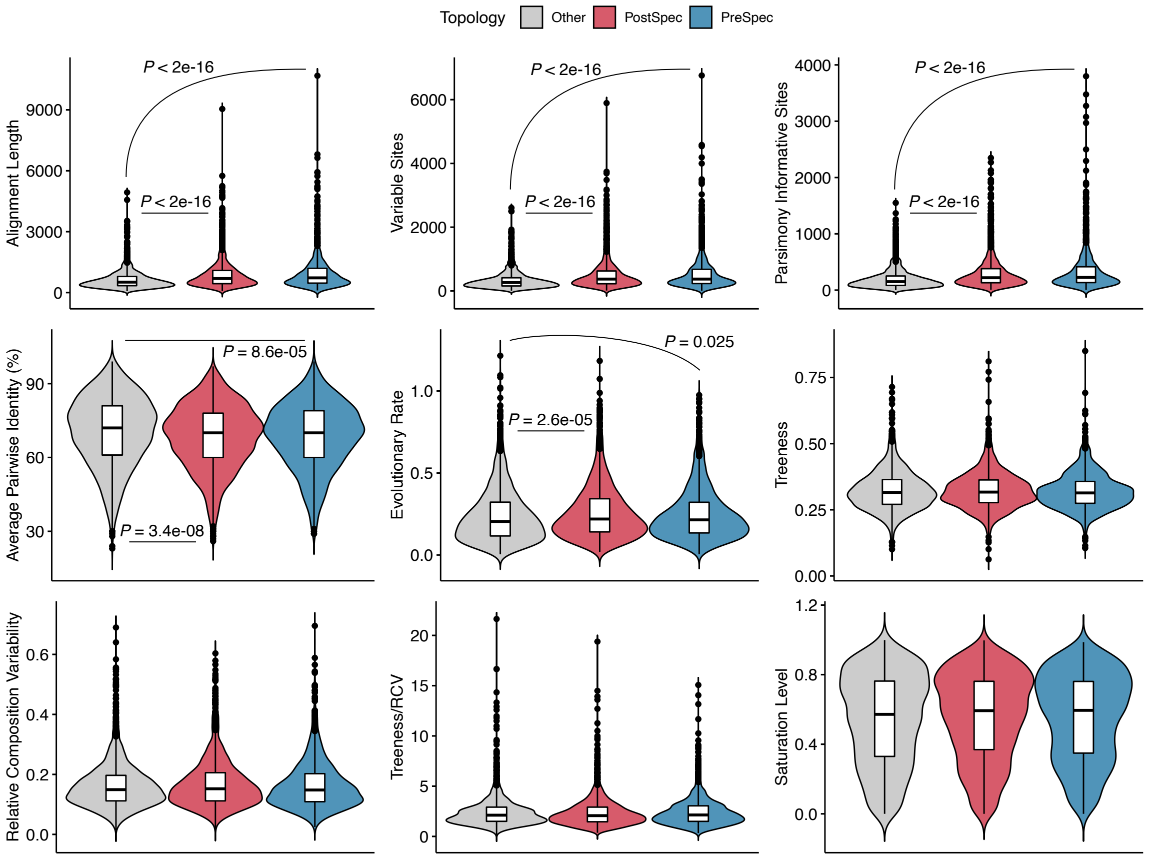


**Figure S1.** Full violin and boxplots plots, as well as significant p-values for summary statistics comparing the three topology categories reported in Figure 3E.

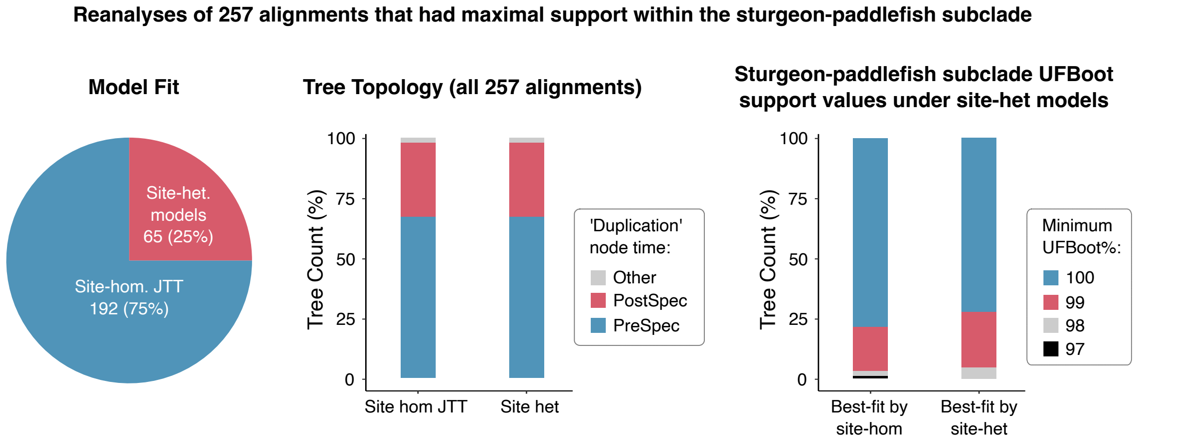


**Figure S2.** Summary statistics for reanalyses with site-heterogeneous mixture models of the 257 alignments for which maximal support (UFBoot=100%) was recovered for the two branches in the sturgeon-paddlefish subclade. The proportion of alignments better fit by site-heterogeneous models as compared to the site-homogeneous JTT model are shown in a pie chart on the left. The frequency at which each of the three topology categories are recovered when always using the best-fitting site-heterogeneous models (including for the 75% of alignments where site-homogeneous models fit better) as compared to when all alignments are analysed with the site-homogeneous JTT model is shown in the centre. The impact that using site-heterogeneous models has on support values for all 257 alignments is shown on the right, with alignments better fit by site-homogeneous JTT or site-heterogeneous models shown separately.

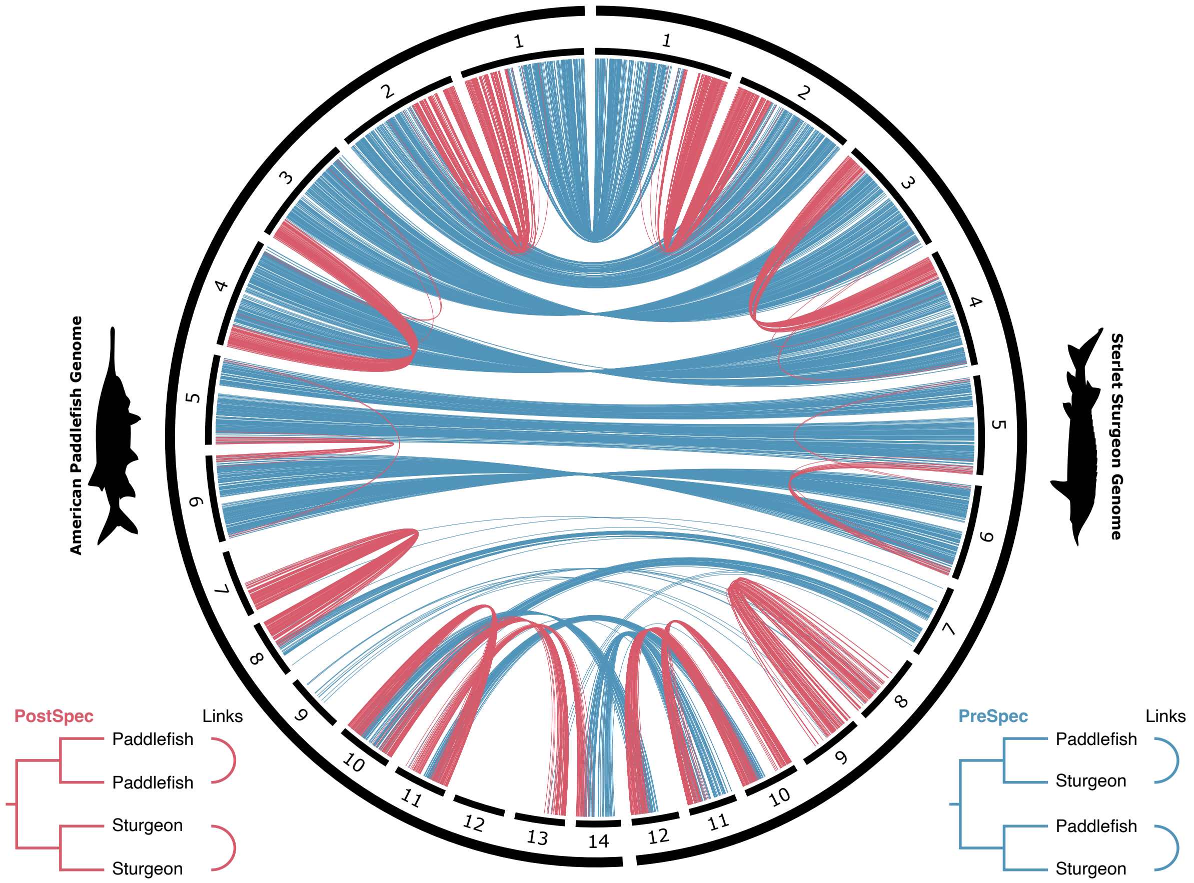


**Figure S3.** Circos plot showing synteny of PreSpec and PostSpec ohnolog pairs links as per Fig. 4C, but for the macrochromosomes >40Mb only.

**Figure S4 (below).** Circos plot showing synteny of PreSpec and PostSpec ohnolog pairs along the sturgeon (left) and paddlefish (right) genomes as per Fig. 4A and Fig. 4B at increasingly stringent UFBoot cut-off% values from top to bottom (UFBoot ≥50%;   UFBoot ≥75%;   UFBoot =100%).

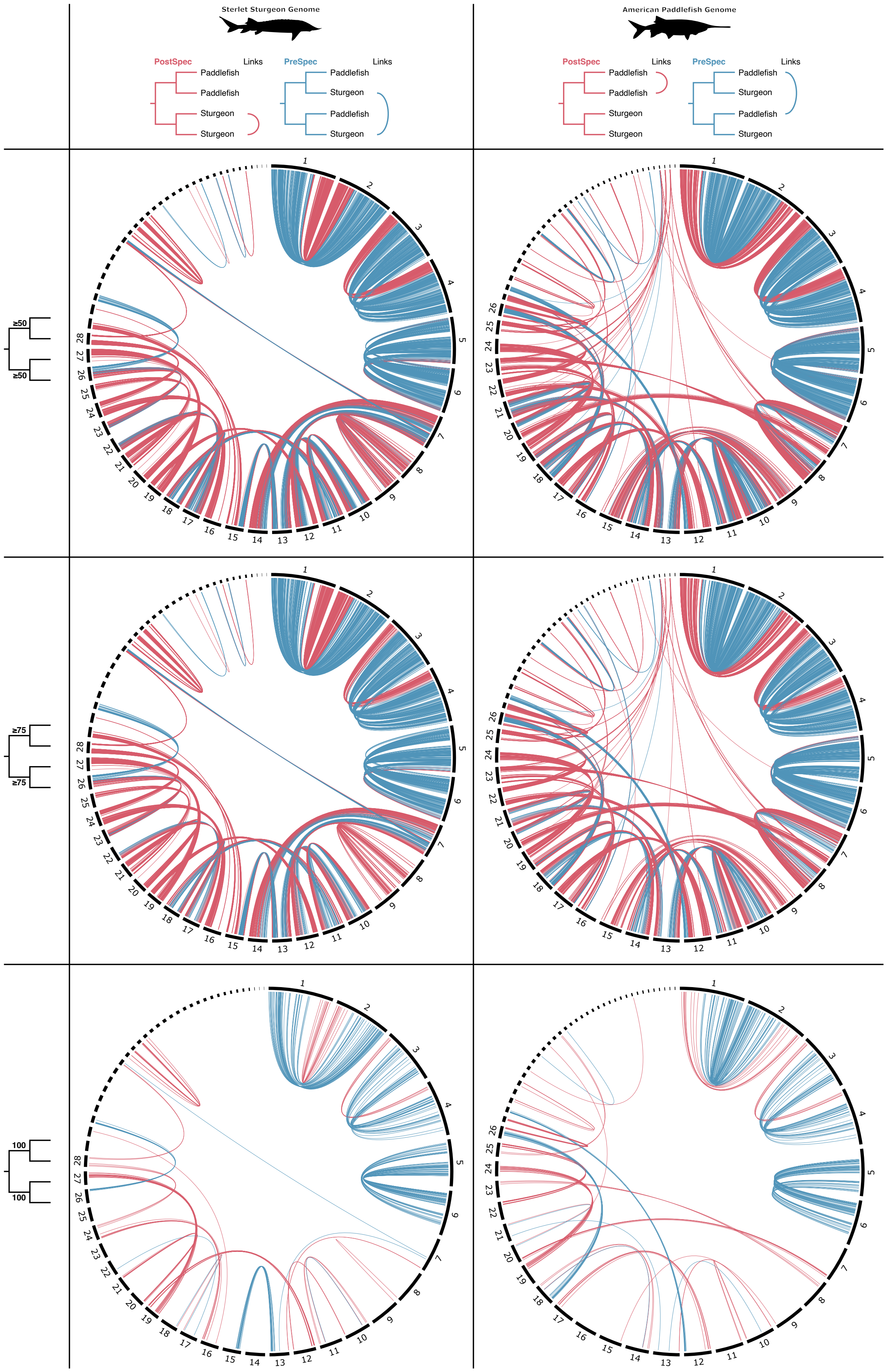

Sterlet Sturgeon Genome
American Paddlefish Genome
PostSpec
Links
Paddlefish
Paddlefish
Sturgeon
Sturgeon
PreSpec
Links
Paddlefish
Sturgeon
Paddlefish
Sturgeon
≥50
≥50
≥75
≥75
100
100

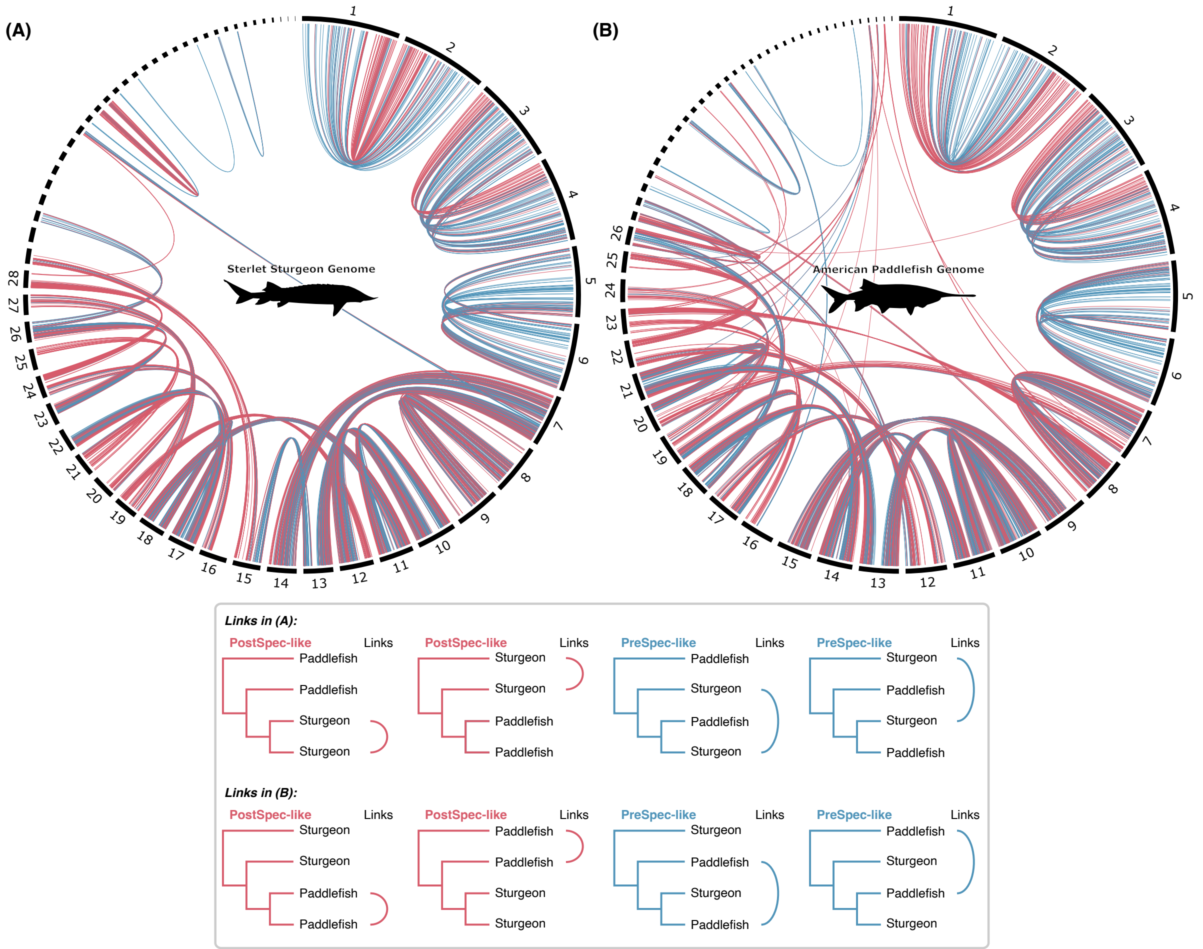


**Figure S5.** Circos plots of synteny patterns of Other topology ‘PreSpec-like’ and ‘PostSpec-like’ recovering ohnolog pairs in the paddlefish and sturgeon genomes. Other details as per Fig. 4A and Fig. 4B.

**Figure S6 (below).** Phylogenomic dating analyses as recovered from all 5 random recodings with mean divergence dates and 95% credibility intervals (blue bar) shown for each node. Runs under the prior are shown on the right.

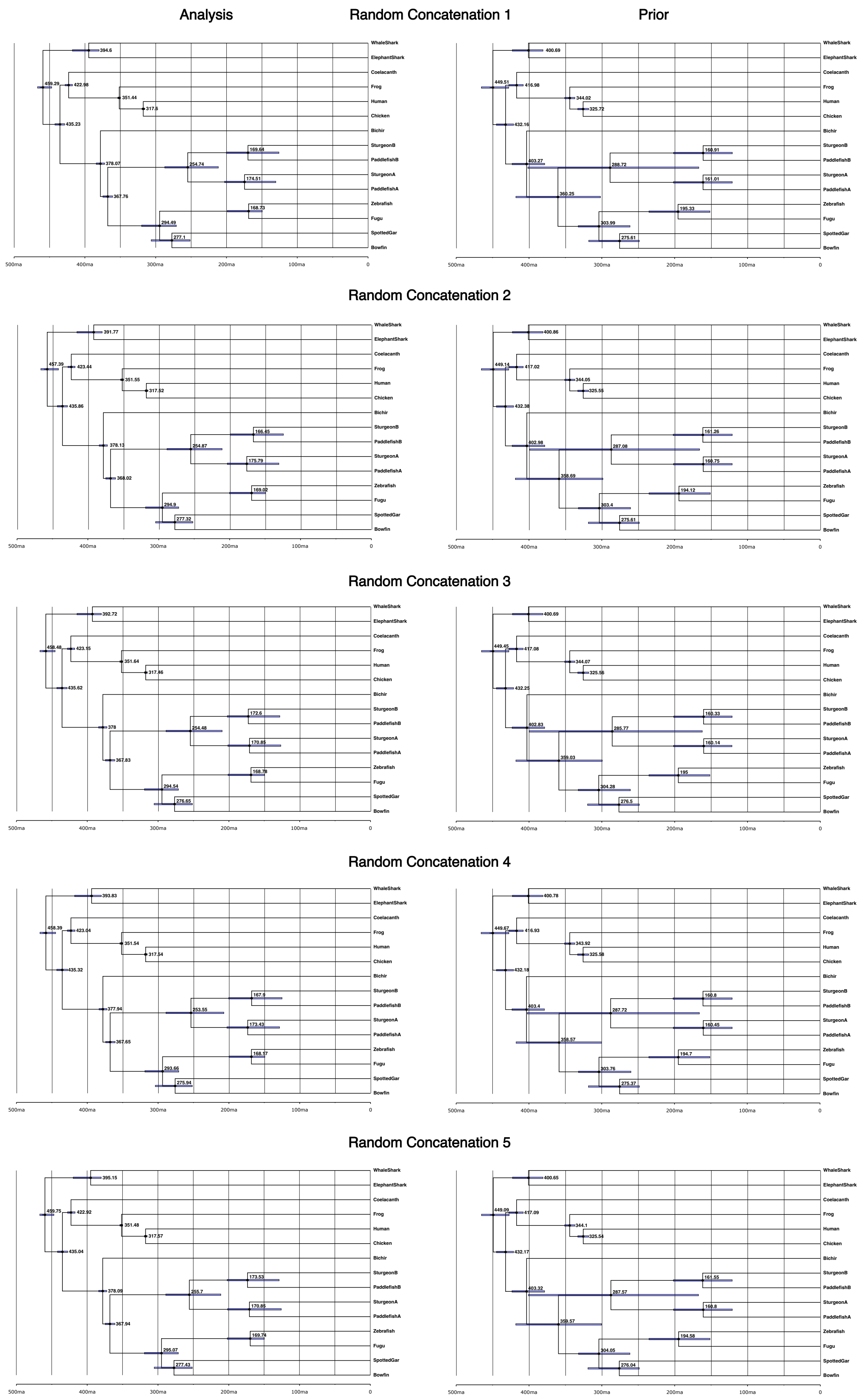
Analysis
Random Concatenation 1
Prior
Random Concatenation 2
Random Concatenation 3
Random Concatenation 4
Random Concatenation 5
WhaleShark
ElephantShark
Coelacanth
Frog
Human
Chicken
Bichir
SturgeonB
PaddlefishB
SturgeonA
PaddlefishA
Zebrafish
Fugu
SpottedGar
Bowfin
500ma
400ma
300ma
200ma
100ma
0
